# HMMSTRTM: A hidden Markov model for local structure prediction in globular and membrane associated proteins

**DOI:** 10.1101/2023.02.08.527695

**Authors:** Tiburon Benavides, Christopher Bystroff

**Affiliations:** Department of Biological Sciences, Rensselaer Polytechnic Institute, Troy NY 12180

## Abstract

**Motivation:** We present HMMSTRTM, a Hidden Markov Model (HMM) that is useful for predicting topology of trans-membrane (TM) proteins. HMMSTRTM provides additional prediction categories of TM regions provided by the PDBTM corpus such as transmembrane beta sheets, coils, and reentrant loops.

**Results:** HMMSTRTM is competitive with existing TM protein topology predictors like TMHMM, it correctly predicts at least half the residues in 96.18% of all transmembrane helices in a cross validation dataset.

**Availability:** Model architecture, source code, and supplementary figures are made available on github: github.com/TiburonB/HMMSTRTM.

## 1. Introduction

Hidden Markov Models (HMMs) have an established track record as predictive tools of protein structure given a protein’s Amino Acid (AA) sequence. Following a popular tutorial of HMMs (Rabiner, 1989), they were applied extensively in computational biology in the 1990’s to find coding regions in DNA (Krogh, et al 1994) to predict protein secondary structure (Asai, 1993), and to describe TM protein topology and predict the presence of TM helices (Krogh, et al 1994). Feed-forward HMMs are used to predict protein sequence families or superfamilies (Finn, et al 2011), and are capable of detecting distant homologs and correctly aligning their sequences.

HMMSTR, (HMMSTR2K) (Bystroff, et al 2000) stood unique in this field as an HMM which could be used to predict dihedral angles, aiding in local structure prediction. HMMSTR2K was constructed by aligning sequence-structure motifs from the I-sites library (Bystroff & Baker, 1998). Positions within the I-sites motifs that co-occurred frequently in the database were merged to form hidden Markov states. A single non-emitting state called a ‘naught state’ was added to connect all C- and N-terminal “dead-end” states, ensuring that the model would return a non-zero probability for any protein sequence. The HMM was then trained on a representative dataset of protein structures and AA profiles using the expectation maximization (EM) procedure (Rabiner, 1989). The trained model was able to predict protein local structure represented as backbone angles; correctly assigning about 69% of all 8-residue segments of all chains in a cross validation subset of the database and about 74% of all 3-state secondary structure descriptors (Q3). HMMSTR2K has since been used to predict contact maps (Bystroff & Shao, 2002), to align remote homologs (Huang & Bystroff, 2006), to recognize remote homologs using a support vector machine (Hou, et al 2004), to recognize good Rosetta protein designs (Schenkelberg & Bystroff, 2015), as input to a protein unfolding pathway prediction algorithm (Ramakrishnan, et al 2012), as a sequence palette for automated protein design (Huang, et al 2015), and as a source of probability-based force fields for C-alpha-only molecular dynamics (Buck & Bystroff, 2009).

First order Markov chains are used to describe a dynamic world state over a time interval and are most easily understood by describing the forward algorithm. In this algorithm, the probability of being at a particular state *j* at time *t* is dependent on the observation of a world state *O* at time *t*, the probability of being in an adjacent connected state *i* at time *t-1*, and the transition probability between states *i* and *j*. HMMs provide an avenue by which unobservable world state information can be predicted based on a sequence of observable information. HMMs used to describe protein structure are not based on time, but rather a position in a sequence. Thus, the information which is predicted by HMMSTR at position *t* in a protein sequence will depend directly on the distribution of states reached at position *t-1*, the descriptive power of each state encountered at position *t*, and the transition probabilities of each state *i* at *t-1* to each state *j* at t. HMMSTR’s states contain several descriptors (emissions) of sequence level information including: an AA profile, secondary structure categories provided by the program DSSP (Frishman; Argos, 1995) Ramachandran (backbone torsion angle) space, and transmembrane categorization provided by PDBTM (Kozma, et al 2012).

An HMM’s predictive capability can be improved stochastically via an expectation maximization (EM) procedure. The EM procedure calculates a *n*x*t* matrix, alpha, via the forward algorithm described above and a similar matrix, beta, which is found by traversing the states backwards, from state *j* to *i* where a directed transition exists *i* → *j*. The backward algorithm begins at time T with equal likelihood to be at all states and ends at time 1. The alpha and beta matrices are then used to calculate other matrices, gamma and zeta, which relay the probability to be at any state *i* at position *t* and the probability to transition from state *i* to state *j* at position *t*, respectively. These matrices can be used to optimize parameters of the model including the probability of transitions between any previously connected states *i* to *j*. The algorithm can remove state transitions entirely, however new transitions will never be added. Analogously, the gamma matrix can be used to weigh state emission predictions via maximizing each state to describe the distribution of observations presenting themselves to the state across all observations in the dataset. For more details see Rabiner 1989.

This paper describes the recent development of HMMSTRTM, a model addition to HMMSTR2K which can aid in the prediction of TM regions within a protein’s sequence, while also offering the local structure predictive capabilities of HMMSTR2K. To accomplish this, significant inspiration was taken from a popular HMM used to predict TM topology, TMHMM (Krogh, 1988).

The topology of TMHMM was defined to mimic the biology of TM proteins. Contained within the model architecture exists sets of states which define globular components of a protein residing on the cytoplasmic and non-cytoplasmic sides of a membrane. Between these globular model regions are a series of connected states which are trained to predict transmembrane helices (TMH). Within these TMH regions is a single “m” state which contains all parameters necessary to determine the optimal length of the TMH (a value constrained between 15 and 35 residues). Five cap states flank the TMH core states. Cap states represent residues which must pass between polar heads of the membrane and are associated with more polar behavior. Two TMH regions exist in TMHMM to create a directional pattern. One region spans the membrane from the inside to the outside and the other which spans from outside to inside. This topology proved fruitful in predicting TMH regions of sequences determined to belong to TM proteins, and since its creation has been used to aid in experimental Protein Structure Determination (PSD) (Tani, et al 2021), and prediction of sequences belonging to TM proteins (Shinzato, et al 2021; Gao, et al 2021).

A mechanism used by TMHMM to preserve model generality and curtail overtraining is the use of ‘tied states’ within disparate regions of the model. Tying a set of states ensures that parameters such as the outgoing transition probabilities or state emissions are kept equivalent within this set of states (Figure 1). For instance, the states within TMHMM’s TMH core are B-tied, meaning that the expected distribution of AA’s is kept constant between all of these states. Likewise, inside cap states are tied together, outside cap states are tied together, and globular inside and outside states are tied together. This last pattern is used to maintain biological knowledge offered by the ‘positive inside rule’ (von Heijne, 1986; Jones, Taylor, & Thornton, 1994; Persson & Argos, 1994; Wallin & von Heijne, 1998) which states that positively charged residues like Arginine and Lysine are an important part of defining cytoplasmic globular loops of TM proteins.

**Figure 1.**
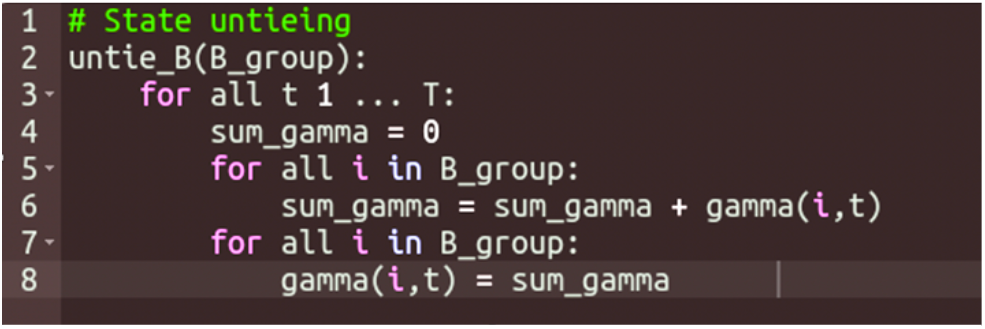
Pseudocode for tied state gamma calculation

## 2. Methods

### HMMSTR2K Topology

During the development of HMMSTRTM, the topology of HMMSTR2K underwent several alterations worth noting. 29 states from HMMSTR2K were identified as redundant and culled from the model greedily by removing states which had the lowest descriptive power across the database. This metric was calculated in each state by summing the EM variable gamma across the database.

A single naught state exists in the model to be used as a master sink for all dead end states in the model. This state ensures no flow is lost in the forward algorithm and the model can efficiently transition from any motif into any adjacent motif. The naught state is well connected, containing 52 incoming transitions and 45 outgoing transitions in the globular model. This state is also non-emitting, meaning it has no predictive value to the model other than to act as an intersection between disparate local structure motifs. Furthermore, this state takes no time step to traverse; meaning if the state is used at time *t*, we will have transitioned from all incoming states at time *t-1* to all outgoing states at time *t*. In the backward algorithm the opposite is true; when the naught state is used at time *t*, we transition from all outgoing naught states at time *t+1* to all incoming states at time *t*. In HMMSTR2K, the naught state has a high prior value, meaning it is used frequently to begin prediction, an alteration was made which disallows starting at the naught state, instead the model’s prior values are trained from the observed probability to be at each state at time *t* = 1. This parameter was initialized so that the model begins at each state with equal probability. (Late in model development, an effort was made to remove the naught state entirely, this vastly increased parameter space and only led to a minor improvement in a derived metric, MDA.)

Naught states can be connected to other naught states, this condition is handled by a recursive function in the forward, backward, and Viterbi algorithms to allow general traversal through linked naught states which do not contain a cycle. Pseudocode is provided which details this recursive naught state traversal (Figure 2). Naught states are used in the HMMSTRTM model to promote a sequential relationship between entering and leaving different regions of the model. For instance, one could expect that the density of motifs encountered after leaving a TM spanning region to be different from the density of motifs encountered in a globular protein.

### HMMSTRTM Topology

This section describes the methods used to create the topology of HMMSTRTM, the model which adds to the predictive capability of HMMSTR2K by accommodating prediction of local structure and TM topology of membrane associated proteins.

HMMSTRTM is built around the original HMMSTR2K topology (Bystroff, 2000) and the PDBTM database (ref). The TM Markov state topologies were constructed to mimic the length distribution and sequence patterns of membrane associated segments of proteins. These designations include four transmembrane regions (TM-regions) which span the membrane in one direction and two that do not cross the membrane, as defined in (PDBTM reference): transmembrane beta-sheet (B), transmembrane coil of unknown structure (C), transmembrane helix (H), transmembrane low-resolution helix (I), membrane integral reentrant loop (L), and membrane interfacial helices (F). Designations are denoted TMx, where x={B,C,H,I,L,F}. TMF states have a route to and from the other TM core regions which skip traversal through cap states. Finally, the TML region contains transitions into and out of globular states on one side of the model.

Data mining strategies, inspiration from existing TM topology predictors like TMHMM, and biophysical intuition were used to construct the TM-region topology of the HMMSTRTM model. To begin, histograms of the periodicity of TM-regions were found to describe the pattern of the lengths of TM regions dependent on the PDBTM-designated character. Following this, an automated procedure constructed Markov chains which mimicked the length distribution of the TM-region. The states of these regions were connected in a fashion similar to TMHMM, where a single state near the center of the TM region ultimately retains the decision for the length of the TM-region. A series of connected cap states flank the TM region. These cap states store patterns in sequence and structural space of the residues which cross over the polar head groups of the membrane. Before training, the states which mark the beginning of the TM region are populated with incoming transitions from all globular states which transition to the globular naught state, and each TM end state was given transitions to each globular state which was transitioned to from the globular naught state. This topology feature highlights the modular nature of working with an HMM defined by local structure motifs; new features can be added as disconnected components from which sequential patterns between disparate local structure motifs can be revealed via EM.

Special consideration was given to TMH and TMB regions of the model. These regions were constructed using an algorithm which establishes the correct consecutive length pattern of the TM region, and appended with rows of states intended to capture the pattern of hydrophilic residues within a TM region. See Figure 3 for details. Other TM topologies were created with similar features, namely a ‘m’ state which arbitrates the length of the TM, a B-tying pattern which maintains important sequence patterns, and cap states which help span the polar head group of membranes. Together four TM topologies, (B,C,H, and I) are used to create a single, directional, TM span between separated globular components of HMMSTRTM, see Figure 4.

**Figure 3.**
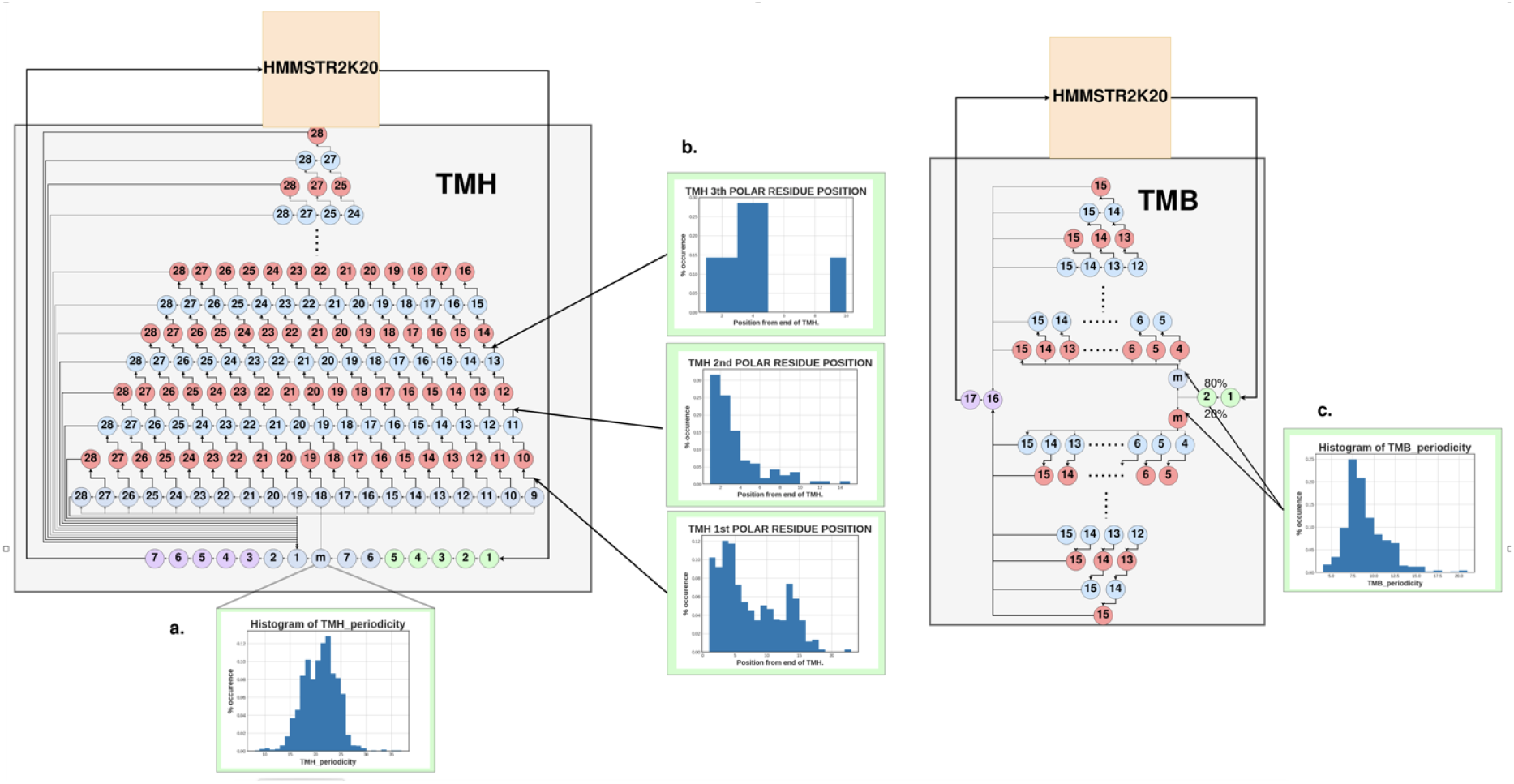
Algorithmically-designed TM region topologies. This figure describes the algorithm-aided of transmembrane helices (TMH) on figure left and transmembrane beta sheets (TMB) extracted from the training database and captured by model parameters; state emissions the TMH length distribution is maintained by setting outgoing transitions of a single of AA polarity across both TMH and TMB topologies in conjunction with state transition ence of polar, or hydrophilic, residues within these protein regions. The line plots in b) the TMH being polar. From this information, we can create a topology which maintains match what is seen in the data. In the TMH region, states have transition probabilities blue states are B-tied and maintain a non-polar AA character. On even layers, red states observation that TMB regions tend to alternate residues of hydrophilic or hydrophobic connected via an alternating pattern of polar and nonpolar B-tied states. The TMB length transition probabilities of two m states. Another state transition controls the frequency nonpolar. Flanking each topology core are TM cap states which are designed to mimic of the membrane. Following topology creation, each TM topology is trained, and states

**Figure 4.**
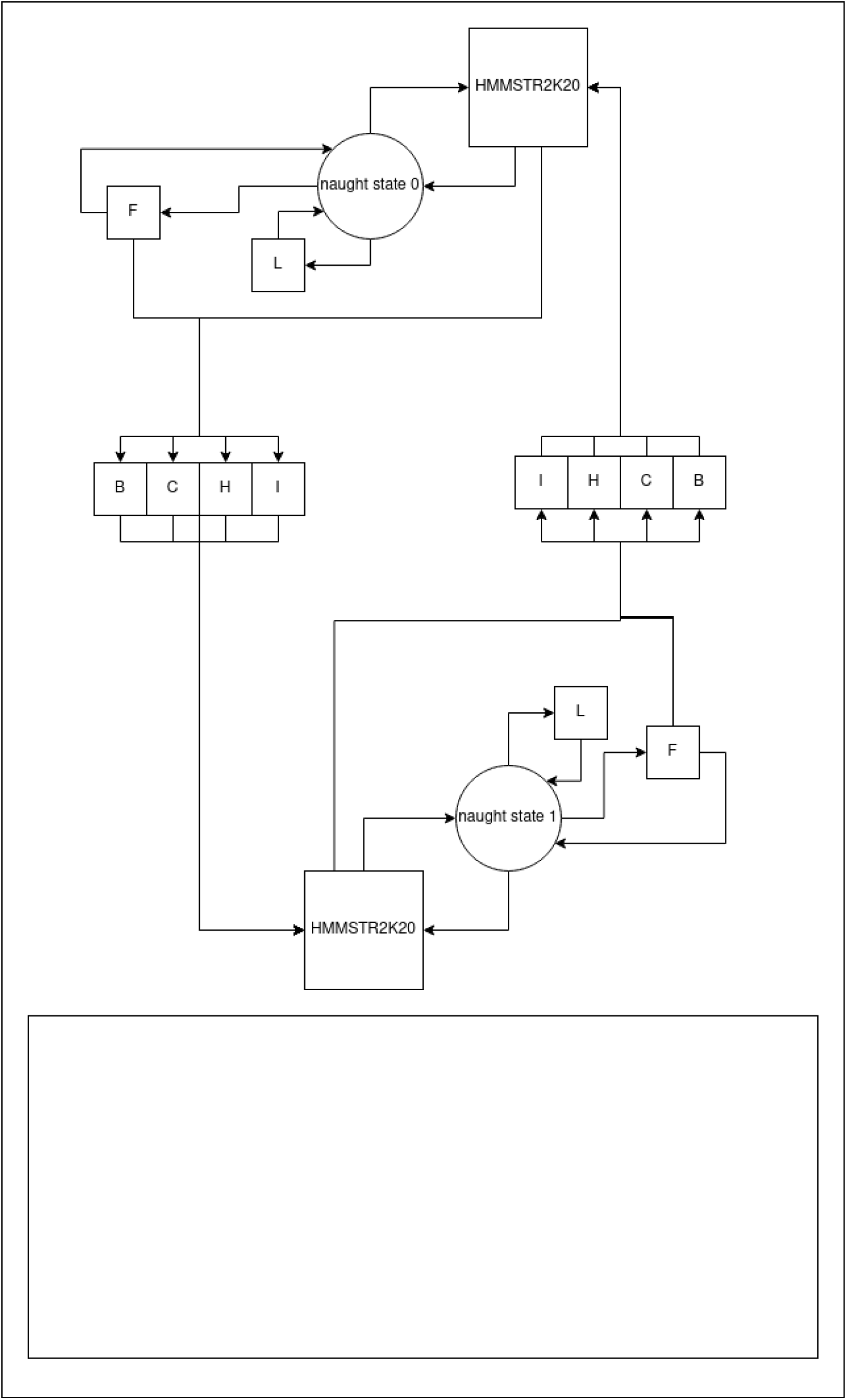
Abstracted the connectivity STRTM model. model STR2K cytoplasmic and loops. A sequence of TM spanning bounds of the diagram are the and relatively complicated transitions between

### Training Data

HMMSTR2K used a training and test set of 691 and 54 protein sequences each. These datasets were pruned for distantly related protein sequences such that no pair of similar sequences would appear in either set or between each set. In the past twenty years, the rate of protein structure determination via experimental methods has followed an exponential curve (RCSB Protein Data Bank, 2021). To account for this, a new database was cultivated from a culled set of proteins from Dunbrack 30% ID cutoff (Wang & Dunbrack, 2003), consisting of 3572 and 655 proteins in the training and test set, respectively. Another dataset containing transmembrane proteins was collected from the PDBTM corpus. This training and test set contains 4737 and 970 chains, respectively, and contains 488 and 127 TM protein chains, respectively. Chains were grouped into protein families when any hit in a blast output was found in a different chain’s blast output. Chains belonging to the same protein family were weighted in the database inversely proportional to the number of chains in the family. Chains in these datasets were curated similarly to the HMMSTR2K dataset such that no pair of similar sequences appeared in any of the datasets or between corresponding training and test sets.

While curating these datasets, several exceptions were encountered relating to sequence alignments. For instance, PDBTM will sometimes omit labeling disordered regions. Likewise, the program used to assign secondary structure, DSSP, would omit assigning labels to disordered regions, and often mis-labels non-canonical AAs. Errors such as these were taken into account by generating an AA sequence alongside sequences for secondary structure and TM region, then performing a sequence alignment back to the ground trut AA (fasta) sequence. Gap characters for the TM and DSSP sequences are ‘U’ and ‘-’ respectively. Characters used in the Ramachandran sequence (*r*) originate from those used in HMMSTR2K using a Voronoi separation procedure of *r* space (Bystroff, 2000)(Figure 5). A custom perl script was used to assign values for the *r* sequence per residue by finding the nearest center of a Voronoi cell in *r* space. In regions of disorder, a character ‘?’ was used as a gap character.

**Figure 5.**
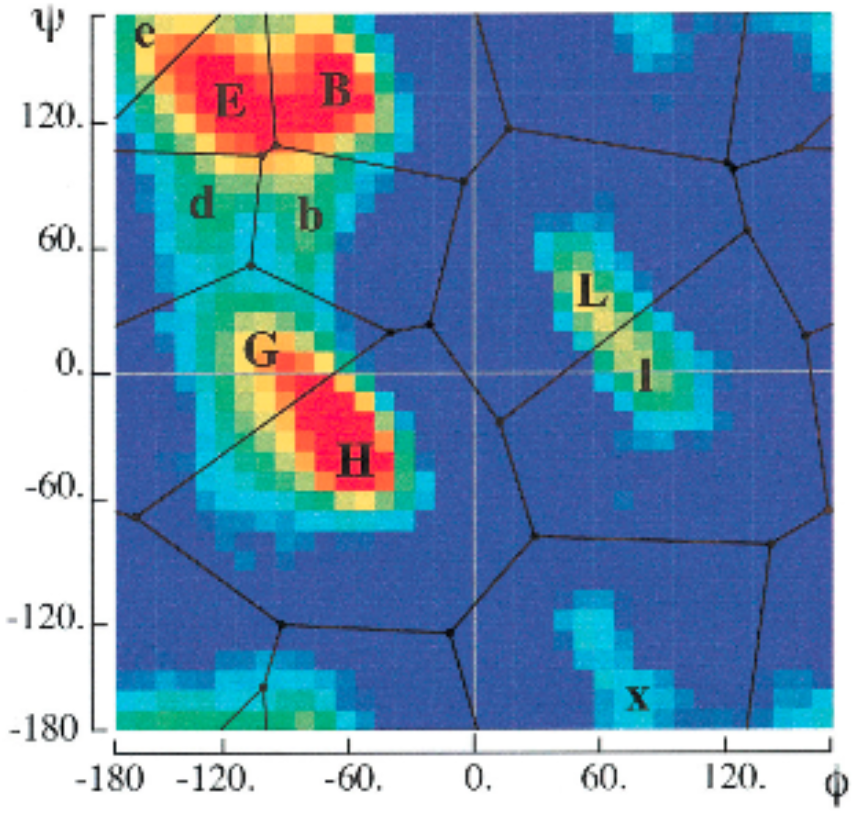
Backbone Angle Regions. This diagram shows regions of Ramachandran (*r*) space seperated by a Voronoi clustering procedure. Areas containing popular dihedral angles are indicated by hotter colors. Not shown is the cis peptide region, ‘c’ (Bystroff, et al 2000).

The PDBTM corpus has several important caveats worth mentioning. Most notably, designated globular components of chains in PDBTM are labeled with the *tm* characters “1” or “2”. However, these labels are not meant to suggest any relationship between ‘intracellular’ or ‘extracellular’. This poses an extra challenge for TM topology prediction since it wasn’t possible to directly train the model on any difference between intracellular or extracellular globular segments. In the final version of the model, these characters were collapsed into a single character “1” which is used to describe any globular region. The *tm* emissions, ‘B’, ‘H’, ‘C’, ‘I’, ‘L’ and ‘F’, constitute the membrane associated regions of the model including B=beta, H=helix, C=coil, I=undefined transmembrane, L=non-crossing re-entrant loop, and ‘F’ = non-entrant interfacial helix. “F” is a helical segment associated with the polar heads of the lipids, which often directly transitions into a TM-spanning region. The final PDBTM character, ‘U’, describes regions of unknown TM-localization, or disordered regions of the protein. While training, if the model encounters a ‘U’ emission, any impact of the *tm* sequence at this time point on the forward-backward matrices is ignored; thus we rely only on the AA profile and *r* character at these positions. During the update step, a ‘U’ emission is counted as a globular region: ‘1’.

### Training

Training was performed on HMMSTR2K with the new dataset. Training used AA profiles and *r* sequences as input provided from the training database previously described. Training via expectation-maximization (Rabiner, 1989) is mathematically guaranteed to increase the predictive capability of the model deterministically to a local maximum. Training for 50 iterations on the dataset resulted in improvement of our metrics and derived metrics when evaluated on equivalent input. However, as most use-cases of HMMSTR will use only an AA sequence or profile as input, it is important to evaluate using this input configuration. Training on AA and *r* input decreases accuracy of successive iterations when evaluated solely on AA profile input. Many attempts were made to circumnavigate this training-evaluation pipeline shortcoming. An obvious solution to this dilemma may appear to train on AA input alone, however this would neglect important local structure information available to the model in the form of the *r* sequence.

HMMSTR’s objective function for expectation-maximization is the probability of the training dataset given the current model, calculated using the forward algorithm (get_alpha in Figure 1). For each possible state pathway through the model, the transitions are multiplied by the emissions, and then the probability of all pathways are summed. The emissions in our case consisted of the state-specific and position-specific AA and *r* likelihood ratios, as shown in Eq 1.

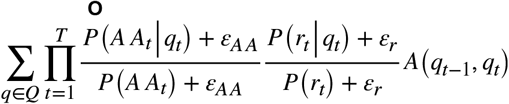

where AA_t_ and *r*_*t*_ are the AA and *r* profiles, ε_AA_ and ε_r_ are the *epsilon* values described below, and q∈*Q*={*q*_*1*_,*q*_*2*_,…,*q*_*T*_} are all state pathways of length T, a *q*_0_ is the *start/end* state.. The training data set sequences are treated as one long sequence with requisite visits to the non-emitting *start/end* state after each C-terminal residue. The objective function **O** is maximized for the training set by iteratively setting the emissions and transitions to their respective expectation values, calculated using *gamma* and *zeta* values from the forward-backward algorithm as described in (Rabiner, 1989). (Eddy, 1998) have shown HMM predictions of protein structural information can be improved greatly by providing an AA profile generated by Multiple Sequence Alignment (MSA) of related sequences revealed by BLAST (Alshul, 1991). HMMs such as this use a cumulative log-odds scoring convention of a given AA profile in the objective function (Barrett, et al 1997). This requires establishing a null model which estimates the ground truth frequencies of each amino acid. To remain consistent with HMMSTR2K, the null model of this study used the background frequencies of each amino acid as observed in our training database. Less common emissions per sequence category are more influential to the log-odds ratio returned from the function. For example, a true positive for a less common AA such as Tryptophan will increment the AA log-odds ratio more than a true positive for a common AA like Alanine. Similarly, incorrectly predicted values will decrease the log-odds ratio for the given emission. Because our AA input takes the form of a *20*t* profile instead of a *t* length sequence, the boundary for what is considered a true positive and a true negative is controlled by the null model, and an additional hyperparameter *epsilon* (akin to a “pseudocount”) which sets a constant minimum value for a zero probability mismatch between any observation and the ground truth per emission category.

The value of *epsilon* is most easily interpreted as a means by which we can control the value of a true negative. With an *epsilon* of 0.001, a ground truth *r* character of ‘L’ and a state with 0% ‘L’ character, we will return a log-odds ratio for the rama emission of -3, which is equivalent to log(*epsilon*). This value can be customized per emission such that the *epsilon* for the AA emission can be different from that for the *r* emission. When the *epsilon* for AA is lower than that of *r*, then a true negative for AA will be more penalizing to the resulting value of the objective function than a true negative for *r*. Experimentation with these values was a balancing act and there is no known way to optimize these values for the task at hand. For the TM models, the *epsilon* for the TM emission was set extremely low (1e-8) so as to greatly discourage the model from mislabeling TM regions as globular and vice versa.

### Structure Prediction

Predictions were performed by finding the gamma matrix of the input sequence on the mode through the forward-backward algorithm. Following this, a sequence profile is constructed by weighting the emissions of every state in the model per time point by the corresponding gamma value. Then a procedure similar to the voting procedure described by (Bystroff, 2000) was carried out to arrive at the final prediction sequence per category. Alternatively, Viterbi could be used to find the most optimal single path through the model which describes the input data. Viterbi performed worse than *gamma* weighted predictions which is consistent with the findings of HMMSTR2K. However, Viterbi did slightly outperform gamma-weighting when predicting TM topology.

Following acquisition of a model, the confusion matrices per emission can be ‘balanced’ by applying scalar multipliers to each character in each prediction category. An automated procedure is performed on a model we wish to evaluate which balances the confusion matrices for SS and *r* emissions. This procedure optimizes a value from the resulting confusion matrix, log(#actual / #predicted), to be equal to 1 for each character. This procedure enhanced the predictive capability of the model by limiting overprediction of the most common character, which is a common problem in machine learning when the distribution of emitted classes is skewed (Wang et al, 2019). The matrices resulting from this procedure give better accuracies than the unbalanced matrix, and also serve to point out flaws in our prediction, e.g. less common dihedral angle predictions like ‘L/l’ are more likely to belong to falsely predicted as ‘E/e/B/b/d’.

TM topology prediction takes place in a post-processing algorithm which assumes the probability of being in a TM-region is proportional to the relative amount of flow going through states in TM-regions at time *t*. Furthermore, we make the assumption that the positively charged globular region effectively simulates a cytoplasmic environment, thus the P(INSIDE) is equal to the relative amount of flow going through these states at time t. Lastly, P(OUTSIDE) is equal to the relative amount of flow going through all other globular states in the model at time t.

### TOPOLOGY VISUALIZATION

HMMSTR has a complicated topology with 252 nodes that exist in connected components of 3 - 16 states which can at any point branch to a different connected component. The topology can be viewed by the program Graphviz (Koutsofios et al, 1999; Gasner & Emden, 2009) by plotting each state and each transition. In Figure 6, the states are color and shape-coordinated to match their most commonly emitted character. For example, states which are more likely to emit ‘H’ than any other *r* emission will be triangles, whereas states which emit any of the *r* emissions often associated with beta sheets will be squares. Amino acids are represented by the inner color of each state, with prolines represented as pink, glycines as green, nonpolar residues as gray and positively or negatively charged residues as red or blue, respectively. Lastly, the TM emission character is visualized by a border color of each state. Globular states have a blue border, TM helices are purple, and so on. Below is a graphviz diagram of the globular model, HMMSTR2K20. Graphviz visualizations proved a potent tool for model debugging of both HMM-STR2K20 and HMMSTRTM.

**Figure 6.**
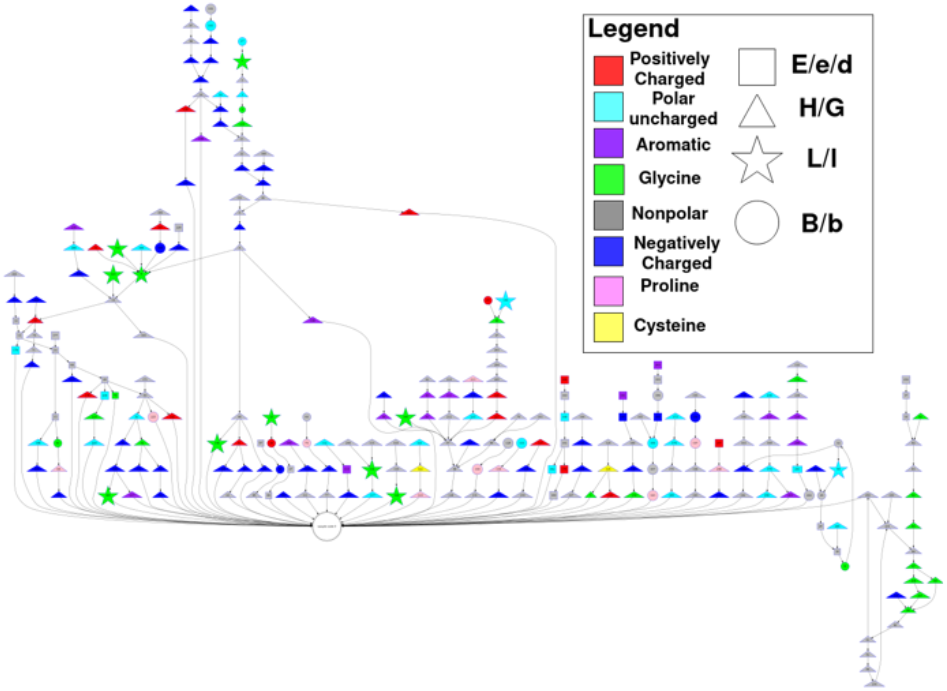
Graphviz topology of HMMSTR2K20. Each state in the model is displayed, with the most prevalent feature of Amino Acid and Ramachandran emissions displayed by color and shape. A well-connected white state near the bottom is the single ‘naught state’ of the globular model, HMMSTR2K20. Transitions below a cutoff value of 10% are excluded from visualization.

## 3. Results

HMMSTRTM is competitive with TMH prediction of TMH-MM (Krogh, 1994), correctly predicting at least half of the TM characters of 96.18% of TM helices across our test set, compared to TMHMM’s performance of 92.28% on the same dataset. Additionally, HMMSTRTM predicts 85.23% of all types TM spanning regions by correctly labeling at least one residue within the region. HMMSTRTM’s overall accuracy for the TM emission is 98.15% across the dataset. Figures 7 and 8 show a typical case.

**Figure 7.**
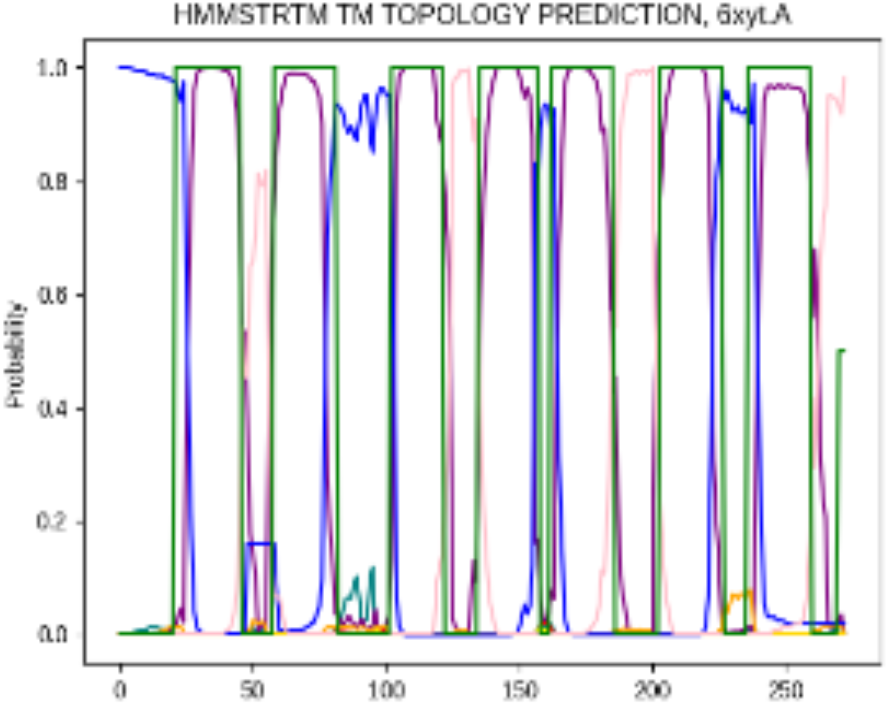
TM Topology prediction of the multi-domain membrane protein, 6XYT.A. Each line in this plot represents the probability of a different TM character. Blue represents cytoplasmic globular, pink is non cytoplasmic globular, purple is transmembrane helix, and green dictates the ground-truth positions of transmembrane helices.

**Figure 8.**
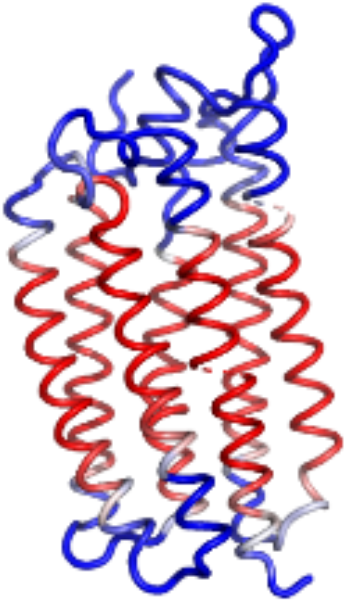
TM region prediction confidence on the TM protein 6xyt.A. Regions in red mark high confidence of residue existing in a transmembrane helix, areas in blue mark high confidence of a residue existing in a globular component of the structure. The last TMH of this chain contains hydrophilic residues within the helix core, however HMMSTRTM is still able to reliably predict

The prediction results were better for alpha helical proteins than for predominantly beta proteins. Upon inspection of beta proteins, the false predictions of backbone angles were found in solvent exposed regions and regions rich in indels in the MSA (Figure 9). A reason for this could be that exposed and evolutionarily variable segments of the protein are last to fold, having the least hydrophobic surface area. Surface segments are first to unfold in the unfolding pathways generated by GEOFOLD (Ramakrishnan, 2021) and are the locations of low phi values (Fowler, et al 2001). The sites of indels in general are places where local structural stability is not as important for folding. The hypothesis therefore is that many surface segments will be incorrectly predicted because they are not in their locally lowest energy conformation. A sequence that prefers helix will be predicted as helix, but if it is a late-folding segment and the structural state of the late-stage folding intermediate prevents the segment from forming helix, then it will adopt a different structure. No three-dimensional structural constraints are encoded within HMMSTRTM.

**Figure 9.**
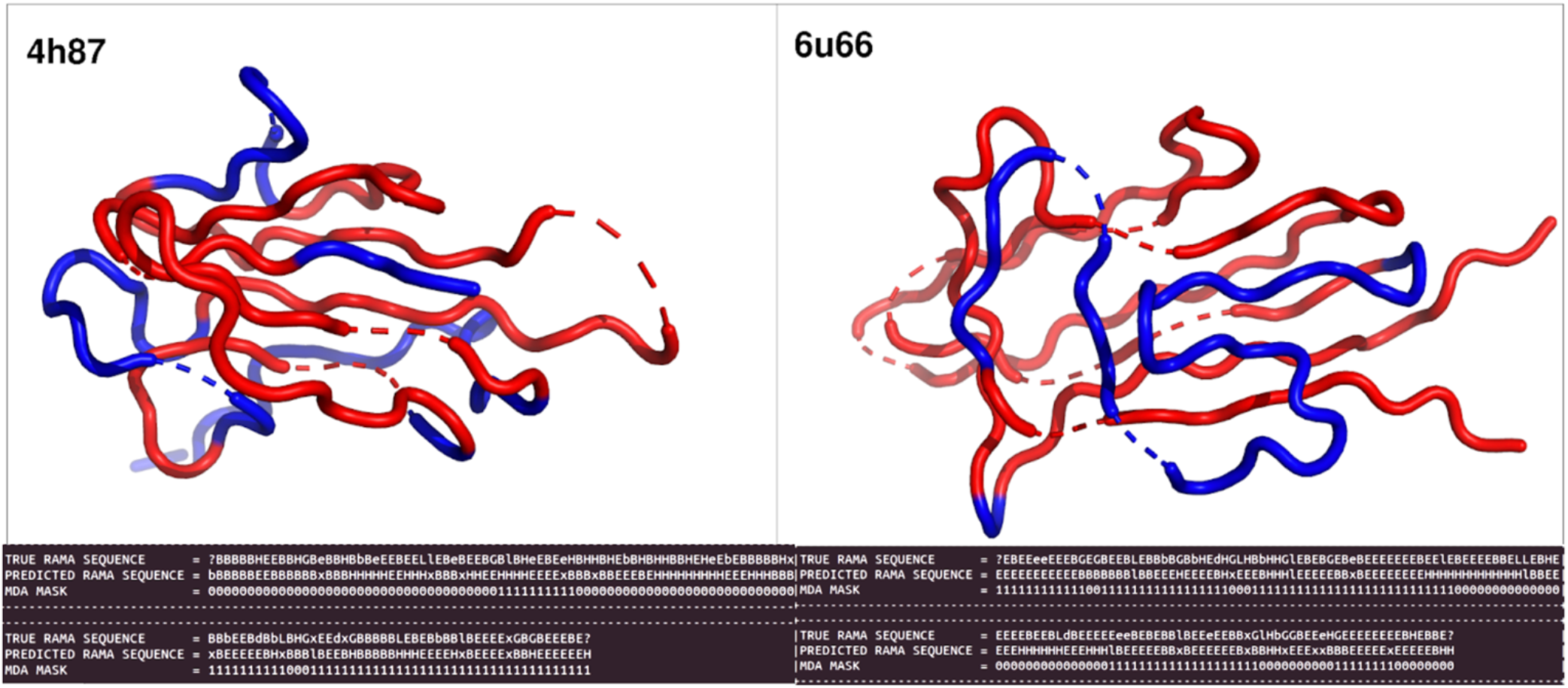
Results of MDA mask on two proteins, PDB IDs: 4h87 and 6u66. These proteins are both primarily comprised of the Beta-Sheet secondary structure. The backbone ribbon structures presented are color coordinated such that red areas were assigned a true value for their MDA mask and blue areas are false. As was common among beta proteins, areas which were commonly assigned false MDA values were those which were likely to be late-folding intermediates of the protein; lacking local sequence support for their local conformation. Below each structure are three sequences relating to the respective protein’s true *r* sequence (RAMA), the predicted *r* sequence, and the boolean sequence resulting from the MDA masking protocol (Bystroff, 2000).

Maximum Dihedral Angle difference (MDA) over an 8-residue window is a derived metric used to determine the accuracy of our backbone angle emissions. If all of the predicted dihedral angles of the 8-mer are all within 120 degrees of the ground truth value then all 8 of these residues are set to true. The first phi angle and last psi and omega angles are omitted when taking the maximum. If a given 8-mer satisfies the condition of MDA it has been shown that the resulting backbone structure will be within 1.0Å RMSD of the actual structure (Bystroff & Baker, 1998). However, if even a single angle within the 8-mer falls outside of the 120 degree boundary, the resulting predicted backbone structure could be quite different from the ground truth configuration, averaging 2.5Å RMSD. HMMSTR2K reported a MDA across a 2000 dataset of ∼69%. This value was recalculated with the untrained model on the new training and test dataset and found to be ∼67%. After training, the value increased slightly to 67.43%.

Confusion matrices are provided to assess the prediction accuracy for residues of each of the Ramachandran regions *r* (Table 3) and to demonstrate the balance between false positives and false negatives. These matrices were calculated across our training and test set to reveal the extent to which the model is overtrained, and help dispel model biases. The most common character per emission category tends to be overpredicted across both datasets, even following confusion matrix reweighting. Another metric assessed model confidence by collecting an ROC value for TM predictions. This value is .714 for the training set and .703 for the test set. Table 4 presents a direct comparison between HMMSTRTM and TMHMM in prediction of TM helix.

**Table 1:**
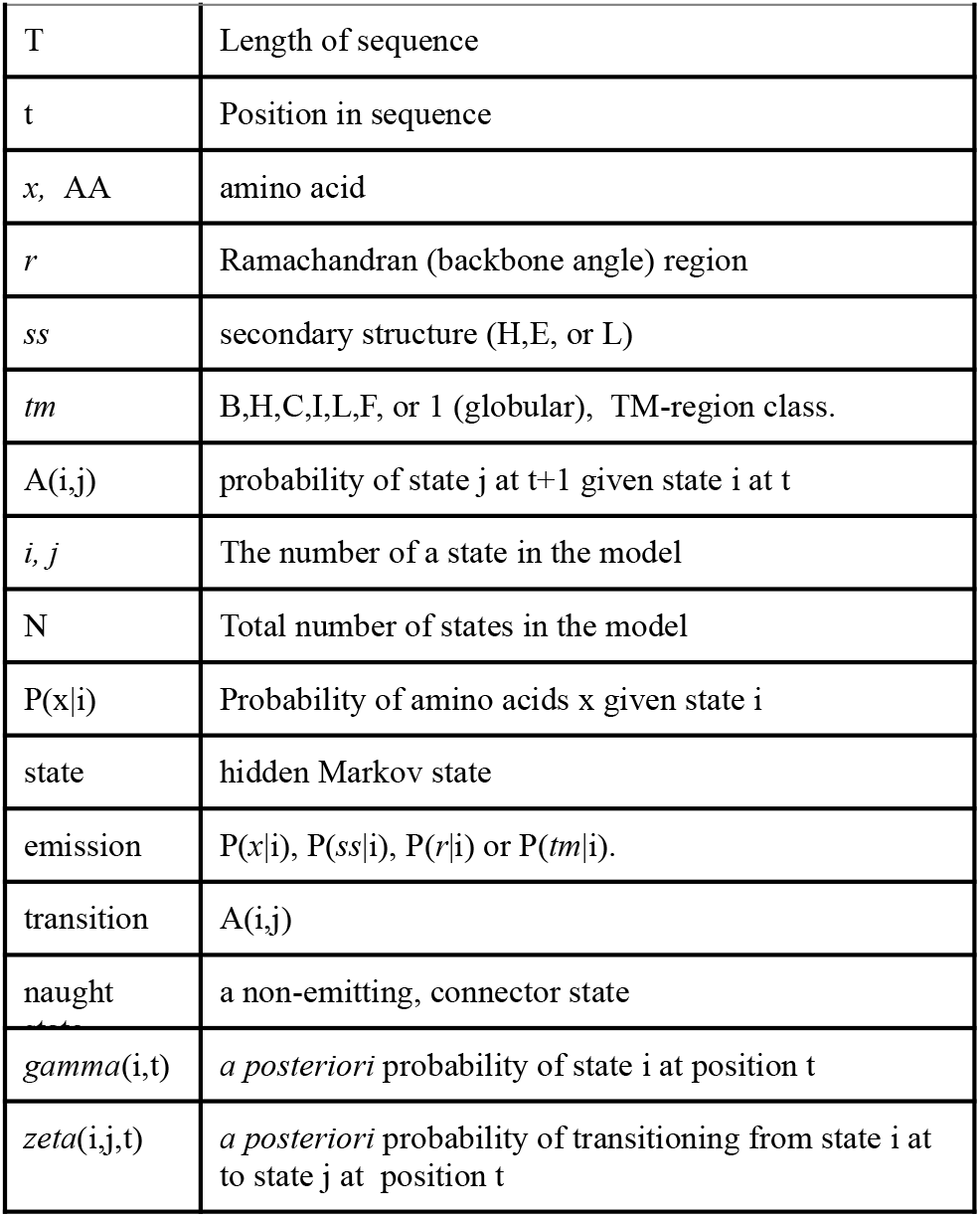
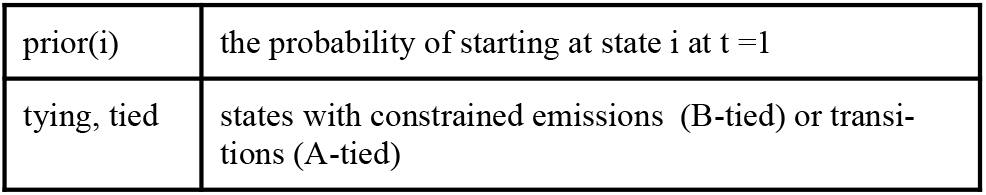
Variable names and terms.

|  |  |
| --- | --- |
| T | Length of sequence |
| t | Position in sequence |
| x, AA | amino acid |
| r | Ramachandran (backbone angle) region |
| ss | secondary structure (H,E, or L) |
| tm | B,H,C,I,L,F, or 1 (globular), TM-region class. |
| $A(i,j)$ | probability of state j at t+1 given state i at t |
| $i, j$ | The number of a state in the model |
| N | Total number of states in the model |
| $P(x\vert i)$ | Probability of amino acids x given state i |
| state | hidden Markov state |
| emission | $P(x\vert i)$, $P(ss\vert i)$, $P(r\vert i)$ or $P(tm\vert i)$. |
| transition | $A(i,j)$ |
| naught | a non-emitting, connector state |
| $\gamma(i,t)$ | *a posteriori* probability of state i at position t |
| $\zeta(i,j,t)$ | *a posteriori* probability of transitioning from state i at t to state j at position t |
| prior(i) | the probability of starting at state i at t=1 |
| tying, tied | states with constrained emissions (B-tied) or transitions (A-tied) |

**Table 2.**
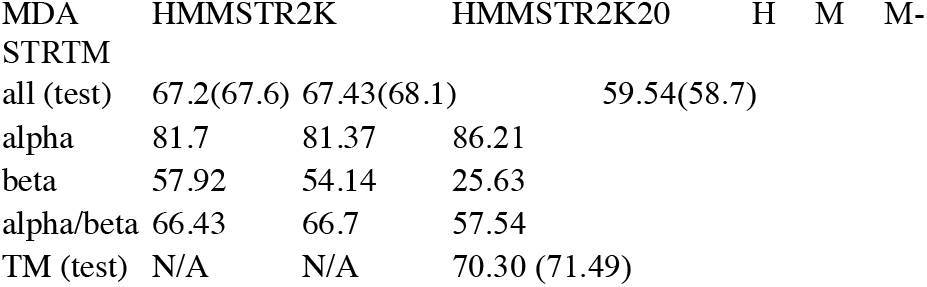
MDA accuracies of the trained and untrained globular model, HMMSTR2K and HMMSTR2K20, and the trained transmembrane model, HMMSTRTM. HMMSTRTM generally overpredicts *r* emissions associated with helices causing the MDA accuracy of the TM model to be much higher in datasets saturated with this secondary structure, and conversely much lower in datasets saturated with Beta sheet secondary structures

**Table 3.**
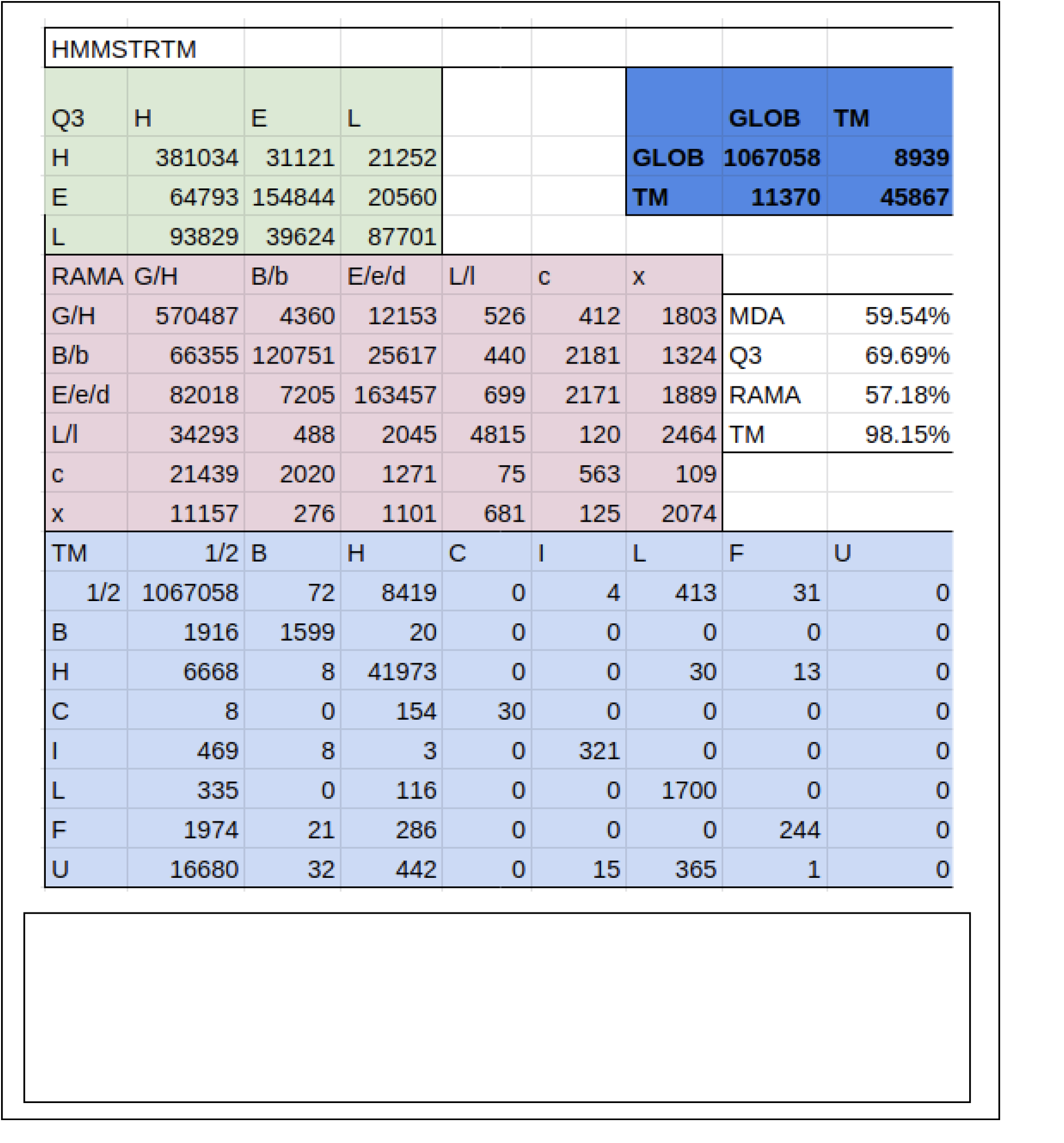

**Table 4.**
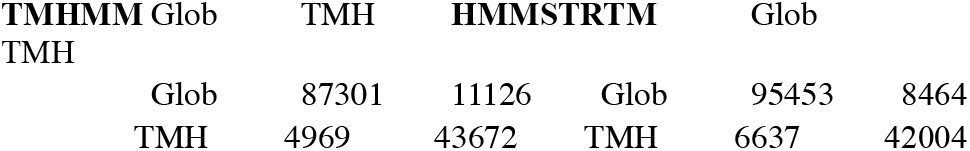
Comparison of TMH prediction between HMMSTRTM and TMHMM. Models compared on a dataset of TM proteins isolated from PDBTM. TMHMM correctly predicts at least one character of 97.7% of TMH; TMHMM correctly predicts at least half of the characters of 92.28% of TM helices. HMMSTRTM correctly predicts at least one character of 85% of all TM regions and 98.23% of TMH regions, and correctly predicts at least half of the characters of 96.18% of TMH.

## 4. Discussion

This study describes the development progress of HMMSTR, a software package which aims to predict the local structure of proteins using sequence motif patterns in the form of a hidden markov model. The new version, HMMSTRTM, also predicts membrane-associated segments of six types: transmembrane alpha helix (H), transmembrane beta sheet (B), transmembrane segment (low-resolution) (I), transmembrane coil (structure unknown) (C), re-entrant integral membrane loops that do not cross the membrane (L), and interfacial segments that sit on, but do not enter, the membrane (F). The new topology blends several existing methods into a single model and highlights the modular nature of Markov Chains.

### Rational design of TM state modules

Great care was taken to ensure the new topology aligned with existing biophysical knowledge of how TM proteins fold and become integrated within a membrane. For example, some trans-membrane helices contain polar side chains, which can occur anywhere in the TM helix but are never adjacent, and TMH are always predominantly nonpolar. Therefore a state topology was constructed that allows isolated polar residues to occur in a predominantly nonpolar sequence segment. This TMH module captures most membrane-spanning helices and excludes globular helical segments.

Similarly, TMB regions are observed to have alternating polar and non-polar side chains reflecting the inside and outside of the beta barrel, respectively. But there are two ways to achieve this, one starting with the polar and the other starting with the non-polar residue. A module that allows either polar or nonpolar to start the TMB segment without loss of the strict sequence pattern was constructed and trained against the database, finding that 80% of TMB start with the non-polar residue and 20% with the polar residue.

Unfortunately, unlike the TMH model, the TMB bears some similarity with portions of the globular model HMMSTR2K20 and therefore produces more false predictions of TMB in sequences that should have been globular beta. The reasons for this may be difficult to remedy, since HMMs are inherently local in their modeling ability. The sequence signals that determine whether a beta sheet protein integrates into the lipid bilayer or not are largely non-local in sequence and involve chaperone proteins such as BAM and TAM in the gram negative bacterial outer membrane (Ranava, et al 2018).

### Reducing model complexity using naught states and tying

The power of modeling using HMMs is enhanced by the inherent ability of this method to be modular. Using naught states and tying were ways to take advantage of HMM modularity while maintaining a low parameters-to-data ratio, as follows.

The addition of a large number of Markov states to a model could potentially lead to overfitting, but techniques were employed here to avoid that pitfall. Specifically, the emission profiles of states were “tied” such that the tied states were updated as a group during EM training. Tying in this way was justified as a model constraint on a case-by-case basis. For the TMH and TMB modules, it was assumed that all transmembrane non-polar positions would have the same intrinsic amino acid preferences, and also that the polar positions would all have the same preferences. This assumption is based on the known interactions of the non-polar side chains with the lipids, which is expected to be position independent.

The HMMSTR2K20 module was duplicated in HMMSTRTM, and this doubled the number of parameters, but the equivalent states in the two models were not tied to reduce the parameter space. By not tying the states, we allowed the model to find sequence pattern differences between cytoplasmic and extracellular globular domains if they exist. There is not clear signal for such differences. But a test for overfitting was performed and the model was found to be not overfit to the training data. Specifically, we determined the accuracy of prediction using the MDA metric on an independent, randomly selected subset of the database which was not used during the EM training, and found that the accuracy was not lower, as it would be if overfitting was happening.

Naught states are non-emitting states that can be used to connect groups of states to other groups. In (Bystroff 2000) a single naught state was used to connect all state sinks to all state sources to assure that the model contained no dead-end states. In HMMSTRTM, naught states were used to connect modules, decreasing the total number of variable state-state transitions from m*n for directly connect modules to m+n for naught state-connected modules, where m is the number of sinks and n is the number of sources. In our experiments, using naught states in this way improved the runtime, an added benefit.

### Uses of HMMSTRTM

This work is part of several efforts in protein structural bioinformatics in the Bystroff lab, and could be useful for any application that needs to assign local structural information to a sequence of unknown structure. We have used HMMSTR2K for several purposes as mentioned in the Introduction. HMMSTRTM will similarly contribute to the applications mentioned. For HMMSUM (Huang & Bystroff, 2006), the new model will provide position-specific amino acid substitution matrices for better remote homolog detection, which will now help identify remote homolog transmembrane proteins. TM regions are expected to have their own specific substitution preferences. For CALF (Buck & Bystroff, 2009), the molecular dynamics program that acts on a minimalist representation protein model, the HMMSTRTM predictions combined with known structures will provide state-dependent, 3D probability fields which will now capture the way TM regions pack in the membrane. For GEOFOLD (Ramakrishnan, et al 2012), the new model will provide assignments of TM regions which will allow us to develop new unfolding moves for those regions, resulting in folding pathways for membrane proteins.

